# Single-cell transcriptomics reveals spaceflight-induced accelerated aging and impaired recovery in the murine bone marrow

**DOI:** 10.1101/2025.11.12.688138

**Authors:** Zhiqun Zhao, Jun Zhang, Guangyu Ji, Zhenzhen Zhou, Yaozong Yang, Fengqi Sun, Haiquan Lu

**Author notes:** These authors contributed equally (Zhiqun Zhao, Jun Zhang). **Corresponding Author** Haiquan Lu, 17922 Jingshi Road, Building 10, Room 204, Jinan, China.

## Abstract

Spaceflight induces physiological changes that resemble accelerated aging, however, how age influences bone marrow response at the cellular level remains poorly understood. Here, we perform single-cell transcriptomic profiling of murine femur and humerus bone marrow from young (12-week-old) and old (29-week-old) mice that underwent a 32-day spaceflight mission followed by a 24-day Earth recovery, with age_-_matched ground controls. Our analysis reveals that, compared with young cohorts, old mice exhibit persistent dysregulation after spaceflight, most prominently in erythroid and B cell lineages. In erythroid cells, old flight mice show pronounced aging signatures, characterized by impaired maturation, inhibited mitophagy, and increased oxidative stress. In B cells, old flight mice show dysregulation associated with failure of the AP-1 stress-response pathway and complete collapse of the intercellular CXCL signaling network. Our findings dissect the age-dependent effects of spaceflight on the bone marrow hematopoietic and immune system at single-cell resolution, and demonstrate that spaceflight imposes a disproportionate burden on the aged hematopoietic system and blunts post-flight recovery. These insights provide candidate pathways and biomarkers for health monitoring and countermeasures in long-duration missions.

## Introduction

As humanity enters a new era of deep-space exploration with planned long-duration missions to the Moon and Mars, understanding and mitigating the associated health risks to astronauts is important. The unique environmental stressors in space, including microgravity and ionizing radiation, induce a series of molecular and physiological changes in organisms that closely resemble the natural aging process on Earth^1–3^. Accumulating evidence indicates that spaceflight accelerates aging-like changes across multiple organ systems. The musculoskeletal system exhibits bone loss and muscle atrophy^4,5^; the immune system shows dysregulation with increased susceptibility to infection and latent virus reactivation^6^; the cardiovascular system shows endothelial dysfunction, vascular remodeling and altered autonomic regulation^7,8^; and the central nervous system exhibits neurovestibular disturbance, neuroinflammation and cognitive effects^9,10^. Among these organ systems, the bone marrow is particularly susceptible, as it plays a critical role in both hematopoiesis and immune regulation^11^. Experiments from the International Space Station (ISS) have revealed complex adaptations within the bone marrow. For instance, astronauts exhibited a marked decrease in vertebral bone marrow adipose tissue 41 days post-flight. This change is temporary and linked to the enormous energy demands of rebuilding red blood cells and bone mass^12^. Notably, younger astronauts showed a greater magnitude of bone marrow adipose tissue (BMAT) modulation^12^, highlighting an age-related difference in recovery dynamics. Besides, impaired lymphopoiesis in mice bone marrow was observed during spaceflight, with granulocyte-macrophage progenitor cells becoming more differentiated^13^.

Despite these insights, much of our current understanding comes from ground-based analog models^14^. Hindlimb unloading studies in both young (2.5-month-old) and aged (18-month-old) mice have shown that mechanical unloading, along with aging, significantly decreases early B cell populations in the bone marrow^15^. This impairment is linked to decreased expression of B cell transcription factors and altered IL-7 signaling^16^. Furthermore, in vitro models of microgravity have demonstrated that bone marrow stromal progenitor cells exhibit impaired proliferation and osteogenic potential, primarily due to downregulation of the Hippo pathway and reduced nuclear activity of key transcription factors Yap1 and Runx2^17^. Spaceflight also impairs hematopoietic stem cell (HSPC) proliferation and differentiation by altering key signaling pathways^18^, such as the Kit-Ras/cAMP-CREB pathway^19^, and by interfering with purine metabolism that negatively affects MAPK signaling^20^. However, these ground-based models cannot fully reflect the complex conditions of actual spaceflight. The fundamental mechanisms, gene regulatory networks, and intercellular communication pathways governing bone marrow hematopoietic and immune responses to actual spaceflight remain poorly understood. To address this gap, we performed a comprehensive single-cell RNA sequencing (scRNA-seq) analysis of bone marrow from young and old mice that underwent a 32-day spaceflight mission followed by a 24-day recovery period on Earth. We aimed to dissect the age-dependent effects on bone marrow recovery. Our results revealed a pronounced acceleration of aging-related signatures in the erythroid cells and identified a profound dysregulation in B cell populations in old flight mice. Our findings suggest that the aged hematopoietic system is particularly vulnerable to spaceflight, leading to persistent dysregulation even after return to Earth. This work provides critical insights into the cellular and molecular pathways that could serve as biomarkers for monitoring astronaut health and guiding countermeasures during prolonged missions.

## Results

### scRNA-seq atlas of bone marrow reveals age-dependent immune dysregulation during the post-spaceflight recovery period

To investigate how age affects the immune system recovery after spaceflight, we analyzed scRNA-seq data of femur and humerus bone marrow samples from OSD-402 and OSD-403 datasets collected in the Rodent Research Reference Mission (RRRM-2)^21^. The experimental design included young (12-week-old) and old (29-week-old) mice exposed to 32 days of spaceflight in the ISS, followed by a 24-day recovery period on Earth before sacrifice. Age-matched ground control cohorts were maintained concurrently.

After stringent quality control and doublet removal, we obtained a high-quality transcriptomic profile of 148,571 cells in total, with eight biological replicates per group: 44,524 cells from 8 ground control old mice (GO), 34,927 cells from 8 flight old mice (FO), 33,937 cells from 8 ground control young mice (GY), and 35,183 cells from 8 flight young mice (FY). We performed dimensionality reduction based on uniform manifold approximation and projection (UMAP) followed by graph-based clustering, and identified nine major cell clusters (Fig. 1a). We manually annotated these clusters according to canonical markers in the CellMarker2.0 database (Fig. 1b, Supplementary Table 1). The identified cell types and their representative marker genes included 44,659 neutrophils (*Retnlg, Ngp, Camp, S100a9, S100a8*), 34,761 B cells (*Ebf1, Pax5, Cd79a, Chst3, Ms4a1*), 34,451 monocytes (*F13a1*, *Fn1, Plcb1, Stxbp6, Arhgef37*), 12,972 NK-T cells (*Skap1, Gm2682, Bcl11b, Cd3g, Camk4*), 5,344 erythroid cells (*Snca, Hba-a2, Hba-a1, Slc25a21, Rhag*), 4,938 plasmacytoid dendritic cells (pDCs) (*Siglech, Grm8, Srgap3, Pacsin1, Slc41a2*), 4,840 adipocytes (*Adgre4, Pparg, Ace, Nr4a1, Cd300e*), and 4,599 macrophages (*C1qb, C1qa, C1qc, Vcam1, Hmox1*).

**Fig. 1.**
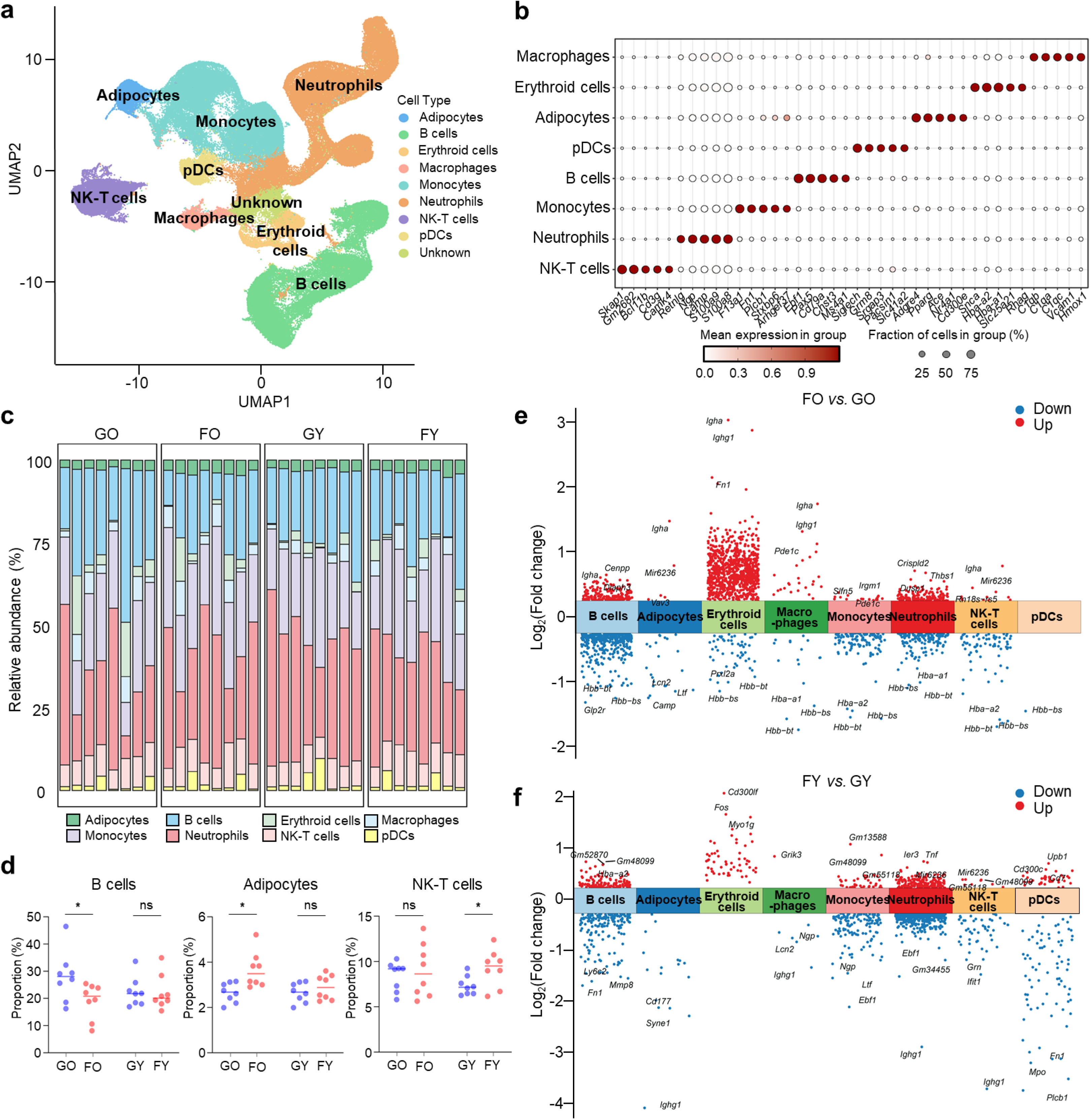
Single-cell landscape of bone marrow samples. **(a)** UMAP plot of single-cell transcriptomes from mouse bone marrow, generated from 32 female mice as part of the Rodent Research Reference Mission (RRRM-2). This dataset comprised samples from four experimental groups: flight young (12-week-old) mice (n=8; FY), flight old (29-week-old) mice (n=8; FO), ground control young mice (n=8; GY), and ground control old mice (n=8; GO). The analysis resolved cells into 9 distinct clusters. **(b)** Dot plot showing the proportional expression levels of typical markers used to annotate each cell type. Dot size represents the percentage of cells expressing each gene; color intensity represents expression levels. **(c)** Proportion of cell subtypes in GO, FO, GY and FY groups. **(d)** Changes in the proportion of B cells, adipocytes and NK-T cells between GO and FO groups, and between GY and FY groups (mean ± SEM; n = 8). \**p* < 0.05. **(e, f)** Differential gene expression comparing FO versus GO samples across cell subtypes in the old group (e) and FY versus GY samples in the young group (f). Red dots represent upregulated genes; blue dot represent downregulated genes.

Compared with mice in GO group, mice in FO group showed a significant decrease in the proportion of B cells (19.11 ± 6.48% in FO versus 28.07 ± 9.23% in GO), and a significant increase in the proportion of adipocytes (3.814 ± 0.66% in FO versus 2.66 ± 0.44% in GO) (Fig. 1c, d), suggesting an accelerated aging process in old mice after spaceflight. In young mice, proportion of NK-T cells increased in FY group compared to GY group (9.25 ± 2.06% in FY versus 7.35 ± 1.04% in GY) (Fig. 1c, d). No significant changes were observed in the proportion of other cell types including neutrophils, monocytes, erythroid cells, macrophages and pDCs (Supplementary Fig. 1a).

We observed that spaceflight had a strong influence on gene expression patterns across cell types. In old mice, spaceflight predominantly influenced gene expression patterns in B cells and erythroid cells, altering the expression of 1,903 and 1,045 genes, respectively (FO versus GO, adjusted *p*-value < 0.05) (Fig. 1e, Supplementary Table 2). In young mice, spaceflight predominantly influenced gene expression patterns in B cells and pDCs, altering the gene expression of 725 and 1,231 genes, respectively (FY versus GY, adjusted *p*-value < 0.05) (Fig. 1f, Supplementary Table 3).

### Dissection and clustering of erythroid cells in the old group

To investigate whether spaceflight accelerates aging in murine bone marrow cells, we calculated the AUCell scores, which quantify the activity of a gene set within each cell, for 11 aging-related gene sets^22^ that we curated from the literature. Among all cell types, erythroid cells in the spaceflight samples showed higher AUCell scores in cellular senescence, stem cell exhaustion, epigenetic alterations and altered intercellular communication compared to ground control samples in both young and old group (Fig. 2a). In addition to erythroid cells, B cells, neutrophils, monocytes, and NK-T cells also demonstrated decreased AUCell scores for several aging-related gene sets (Supplementary Fig. 2a), while adipocytes, macrophages and pDCs showed only slight changes (Supplementary Fig. 2b). In erythroid cells, we identified 582 upregulated genes and 21 downregulated genes in the samples from old group (FO versus GO, adjusted P-value < 0.05, |log_2_(fold change)| > 0.585), but only 42 upregulated genes and 0 downregulated genes in the samples from young group (FY versus GY, adjusted *p*-value < 0.05, |log_2_(fold change)| > 0.585) (Fig. 2b).

**Fig. 2.**
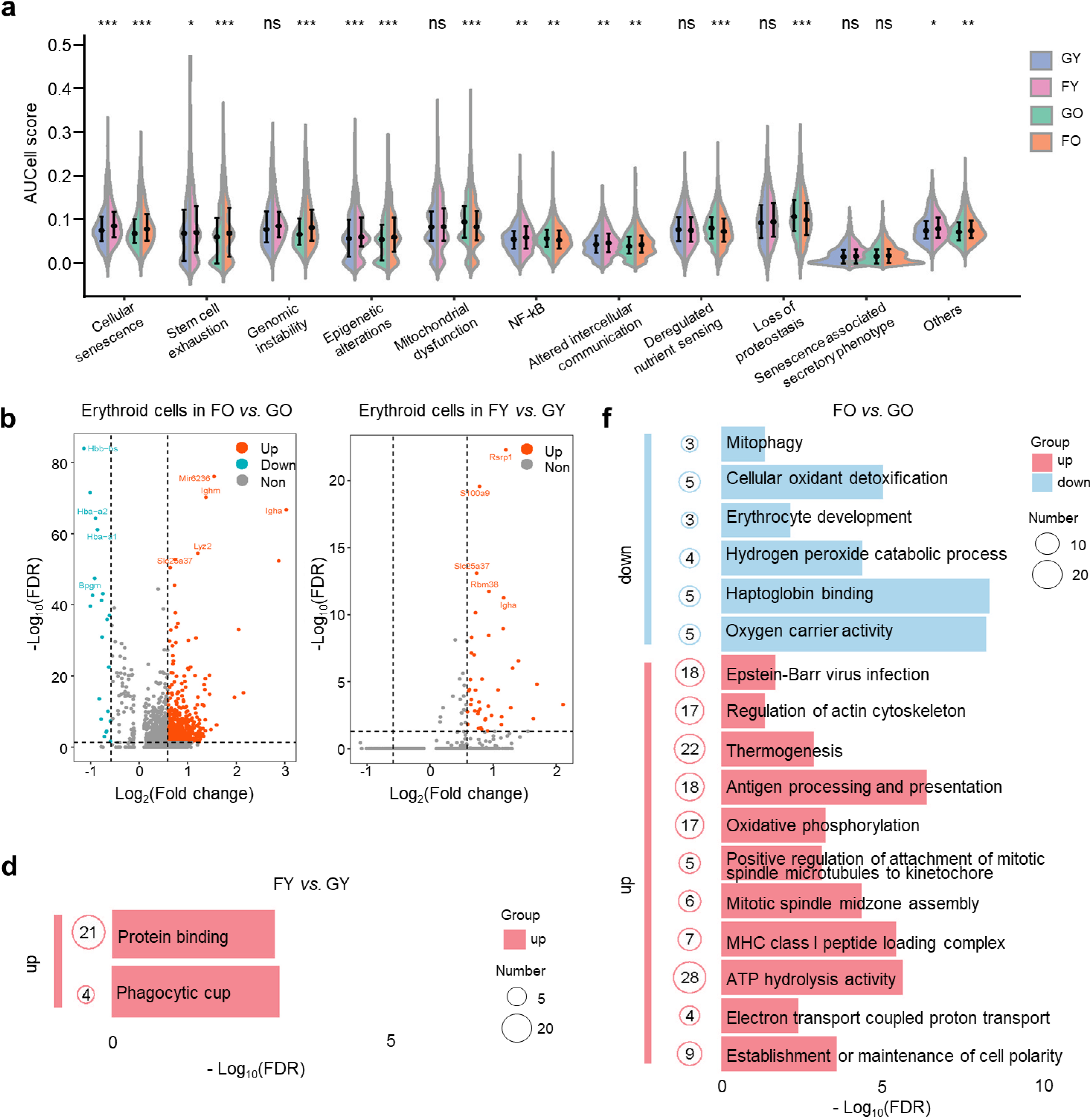
Functional features of erythroid cells in bone marrow samples. **(a)** Comparison of AUCell scores of 11 aging-related gene sets in erythroid cells between GO and FO groups, and between GY and FY groups. \**p* < 0.05, \*\**p* < 0.01, \*\*\**p* < 0.001; ns, not significant. **(b)** Volcano plot showing differentially expressed genes (DEGs) between FO and GO samples in the old group (left panel), and between FY and GY samples in the young group (right panel). Red dots represent upregulated genes, and blue dots represent downregulated genes (adjusted *p* value < 0.05 and |log_2_(fold change)| > 0.585). **(c, d)** Functional enrichment analysis of DEGs in old erythroid cells (c) and young erythroid cells (d).

Gene enrichment analysis of differentially expressed genes (DEGs) in erythroid cells from old mice revealed that upregulated genes were enriched in pathways related to oxidative phosphorylation (mmu00190) and electron transport coupled proton transport (GO:0015990), suggesting enhanced mitochondrial function after spaceflight. Additionally, upregulated genes were enriched in pathways related to regulation of actin cytoskeleton (mmu04810), establishment or maintenance of cell polarity (GO:0007163), positive regulation of attachment of mitotic spindle microtubules to kinetochore (GO:1902425), and mitotic spindle midzone assembly (GO:0051256), indicating changes in erythroid cell morphology and polarity after spaceflight. Downregulated genes were enriched in pathways associated with erythroid cell development, including erythrocyte development (GO:0048821), haptoglobin binding (GO:0031720), and oxygen carrier activity (GO:0005344), indicating impaired erythroid cell development after spaceflight. Notably, the mitophagy (mmu04137) pathway was inhibited, which implies suppressed erythroid development and an increased risk of anemia, because mitochondrial autophagy is essential for the clearance of mitochondria during terminal erythroid differentiation^23^. In addition, pathways related to hydrogen peroxide catabolic process (GO:0042744) and cellular oxidant detoxification (GO:0098869) were downregulated, indicating a decreased capacity to eliminate reactive oxygen species (ROS) and the resulting accumulation of oxidative stress (Fig. 2c, Supplementary Table 4). In contrast, erythroid cells from young mice did not show enrichment of these pathways after spaceflight (Fig. 2d, Supplementary Table 5). These findings collectively indicate that spaceflight exerted long-term impacts on erythroid cells specifically in old mice.

To investigate the heterogeneity among 5,344 erythroid cells, we analyzed their transcriptomic profiles and performed re-clustering analysis. Erythroid cells were categorized into five clusters based on their gene expression profiles (Fig. 3a, b, Supplementary Table 6). In cluster E0, *Bpgm* and *Xpo7* were highly expressed. *Bpgm* is a multi-functional enzyme with a crucial role in metabolic regulation. *Bpgm* converts 1,3-bisphosphoglycerate (1,3-BPG) to 2,3-bisphosphoglycerate (2,3-BPG) through the Rapoport-Luebering shunt, a bypass of the glycolytic pathway, thereby facilitating hemoglobin-oxygen delivery^24^. *Xpo7* is a broad-spectrum nuclear transport receptor^25^ involved in nuclear condensation and enucleation during erythroid nuclear maturation^26^. Therefore, E0 likely represents erythroid cells undergoing terminal differentiation and maturation. Cluster E1 was characterized by high expression of *Stag1*, *Neil3*, and *Pbx3*. *Stag1* is a component of the cohesin complex, which is essential for chromosome segregation during cell division^27^. *Neil3* is a DNA glycosylase involved in DNA repair, which is critical for maintaining genomic integrity in rapidly dividing cells by repairing DNA inter-strand crosslinks^28^ and removing oxidized bases from single-stranded DNA^29^. High expression of these genes suggests that E1 may represent a population of actively proliferating erythroid progenitors. In cluster E2, *Tmem14c* and several ribosomal protein genes (*Rpl41, Rps2, Rps18*) were highly expressed. *Tmem14c* encodes an inner mitochondrial membrane protein required for transporting mitochondrial porphyrins in developing erythroid cells^30^, which facilitates the import of protoporphyrinogen IX into the mitochondrial matrix for heme synthesis and subsequent hemoglobin production^31^. The high expression of ribosomal proteins indicates robust translational activity, which is necessary for synthesizing large amounts of globin chains. This signature suggests that E2 consists of cells engaged in hemoglobinization during erythropoiesis. Cluster E3 showed high expression of genes related to the immune response, including *S100a9*, *Camp*, *Lcn2*, and *Ltf*. *S100a9* is a damage-associated molecular pattern (DAMP) molecule involved in inflammatory responses^32^. *Lcn2* functions as a bacteriostatic agent by interfering with bacterial iron acquisition systems^33^. *Ltf* protects cells from oxidative damage by binding directly to ferric ions with high affinity to prevent the generation of ROS. This expression profile suggests that E3 may represent a subpopulation of erythroid cells mounting a stress or inflammatory response, potentially related to the spaceflight environment^34^.

**Fig. 3.**
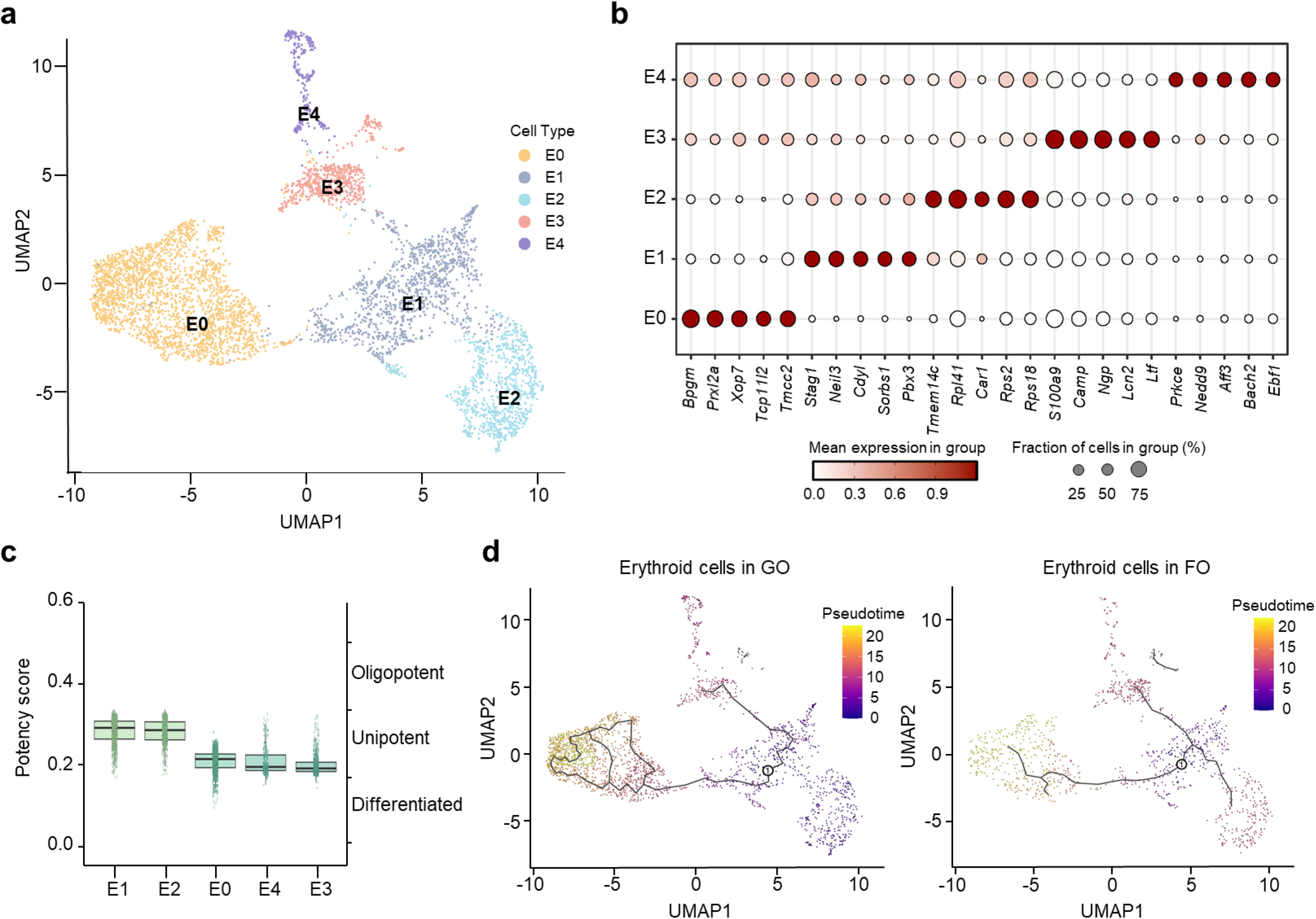
Evolutionary trajectories of old erythroid cells. **(a)** UMAP plot showing 5,344 erythroid cells. **(b)** Dot plot showing the expression of canonical markers used to identify erythroid subtypes. **(c)** CytoTRACE analysis showing differentiation potential of erythroid cell subtypes. **(d)** Trajectory analysis of erythroid cells in GO samples (left panel) and FO samples (right panel). White circles represent the root nodes of the trajectory.

### Evolutionary trajectories of erythroid cells in old mice

We next delineated the developmental trajectory of erythroid cells in old mice. We calculated potency scores by the CytoTRACE algorithm to assess cellular stemness, and found that the E1 subpopulation had the highest score (Fig. 3c), indicating that E1 subpopulation is the least differentiated subpopulations. Therefore, the E1 subpopulation was chosen as the starting point for trajectory analysis. We then performed a pseudotime developmental trajectory analysis and found a significant difference in erythroid development between the GO and FO groups: erythroid cells in the GO group followed a more complete and branched differentiation program while erythroid cells in the FO group exhibited a reduced progression toward terminal differentiation (Fig. 3d).

We next investigated the transcriptional changes associated with transitional states. In the GO group, erythroid subclusters fell into three phases: E1 and E2 in phase 1, E4 and E3 in phase 2, and E0 in phase 3 (Fig. 4a, upper panel). We found that genes in cluster 3 were highly expressed in phase 1 and weakly expressed in other phases. Enrichment analysis of genes in cluster 3 revealed involvement in signaling pathways associated with cell cycle, ribosome, homologous recombination, base excision repair, P53 signaling pathway and ferroptosis. These findings suggested that cells in phase 1 (E1 and E2 subpopulations) are actively engaged in proliferation, DNA repair, and response to cellular damage, indicating that they are in a stage of high activity, repair processes and preparation for differentiation. Genes in cluster 2 were highly expressed in phase 2, and were enriched in pathways associated with B cell receptor signaling pathway, NF-kB signaling pathway, guanyl-nucleotide exchange factor activity and regulation of TOR signaling, suggesting that cells in phase 2 (E4 and E3 subpopulations) are involved in immune response and cellular stress response pathways. Genes in cluster 4 were highly expressed in phase 3, and were enriched in antioxidant activity, suggesting that cells in phase 3 (E0 subpopulation) are engaged in managing oxidative stress and enhancing their antioxidant defense mechanisms (Fig. 4a, lower panel).

**Fig. 4.**
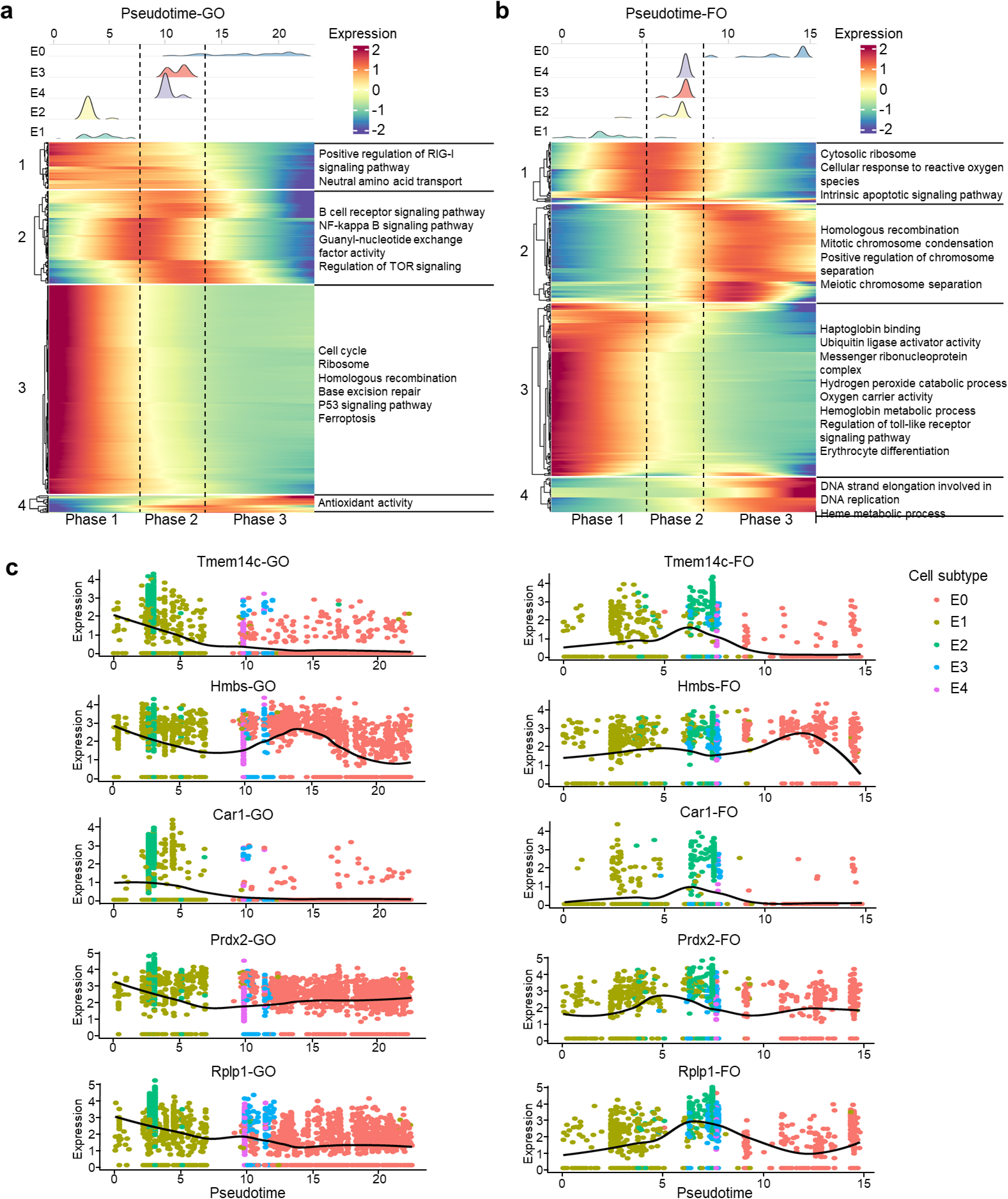
Gene expression dynamics along the evolutionary trajectory of old erythroid cells. (a, **b)** Pseudotime distribution of erythroid cell subtypes during the transition (upper panels) and dynamic gene expression changes along pseudotime (lower panels) in GO samples (a) and FO samples (b). **(c)** Dynamic expression of representative DEGs along pseudotime.

Erythroid cell subclusters in the FO group were also categorized into three phases: only E1 in phase1, E2, E3 and E4 in phase 2, and E0 in phase 3 (Fig. 4b, upper panel). The E2 subcluster in the FO group emerged at a later developmental stage, which is different from that in GO group. We found that genes in cluster 3 were highly expressed in phase 1 and weakly expressed in other phases. Enrichment analysis of genes in cluster 3 revealed involvement in signaling pathways associated with haptoglobin binding, hemoglobin metabolic process, oxygen carrier activity, hydrogen peroxide catabolic process, ubiquitin ligase activator activity, messenger ribonucleoprotein complex and regulation of toll-like receptor signaling pathway, suggesting that cells in phase 1 (E1 subpopulation) were focused on cellular protection and metabolic functions. cluster 1 genes were highly expressed in phase 2 and were significantly enriched in cytosolic ribosome, cellular response to reactive oxygen species and intrinsic apoptotic signaling pathway, suggesting that cells in phase 2 (E2, E3, and E4 subpopulations) were dealing with cellular stress and possibly priming for programmed cell death. Genes in cluster 2 and cluster 4 were highly expressed in phase 3, and were enriched in homologous recombination, mitotic chromosome condensation, meiotic chromosome separation, positive regulation of chromosome separation, DNA strand elongation involved in DNA replication and heme metabolic process, suggesting that cells in phase 3 (E0 subpopulation) were undergoing final stages of cell division (Fig 4b, lower panel).

To identify genes exhibiting distinct expression kinetics along the E2 developmental trajectory, we first identified the intersection of genes between cluster 3 of the GO group (Fig. 4a) and cluster 1 of the FO group (Fig. 4b). These shared genes were then ranked by Moran’s I, a statistic that measures spatial autocorrelation in pseudotime, to select the top five genes most significantly associated with the trajectory (Supplementary Table 7). *Tmem14c, Car1, Prdx2* and *Rplp1* demonstrated different expression kinetics in GO and FO groups. In the GO group, *Tmem14c, Car1, Prdx2* and *Rplp1* expression gradually decreased as pseudotime progressed from early (E1) to late stages (E0), with the highest levels in E1 and E2 subclusters and a decline through the E4 and E3 subclusters (Fig. 4c, left panel). In contrast, in the FO group, *Tmem14c, Car1, Prdx2* and *Rplp1* showed a very low levels at early stage (E1) and gradually increased, with the highest levels in the E2 subcluster and a decline through the E4 and E3 subclusters (Fig. 4c, right panel). Although *Hmbs* exhibited dynamic expression kinetics, it was poorly correlated with the E2 subcluster. Among them, *Tmem14c* is involved in the maturation of erythroid cells. *Rplp1* encodes a ribosomal phosphoprotein component of the 60S subunit, which is essential for embryonic development, brain development, and proper cell proliferation^35^. The expression kinetics of *Tmem14c* and *Rplo1* in the FO group indicated a delayed activation, potentially contributing to slower erythroid maturation in these mice.

Taken together, these data demonstrate that erythroid cells in the GO and FO mice followed distinct activation trajectories. While both groups progressed through three main phases, the spaceflight group showed delayed differentiation. Additionally, erythroid cells in FO group emphasized metabolic and oxidative stress responses in early phases, whereas erythroid cells in GO group focused more on cell proliferation and immune signaling.

### Heterogeneity of B cells between old and young group

B cells (n = 34,761) constituted a major population in the bone marrow samples (Fig. 1a). Using the Seurat package in R, we re-clustered all B cells and revealed seven clustered subpopulations (Fig. 5a). The top 5 marker genes for each B cell subpopulation were shown Fig. 5b (Supplementary Table 8). Notably, B3, B5 and B6 subpopulations demonstrated uniquely high-expressed genes. Cluster B3 was characterized by significantly upregulated *Top2a*, *Tpx2*, and *Nusap1*, all of which are involved in the cell cycle and mitosis^36–38^. Therefore, cluster B3 likely represented a subpopulation of highly proliferative B cells^39^. Cluster B5 represented memory B cells, defined by its high expression of *Fcrl5*. *Fcrl5*^+^ memory B cells have been observed to expand in inflammatory contexts, including chronic infections, autoimmune diseases, aging, and vaccination^40,41^. Cluster B6 demonstrated high expression of *Dntt* and *Vpreb*, which are classic markers of early B cell development^42^.

**Fig. 5.**
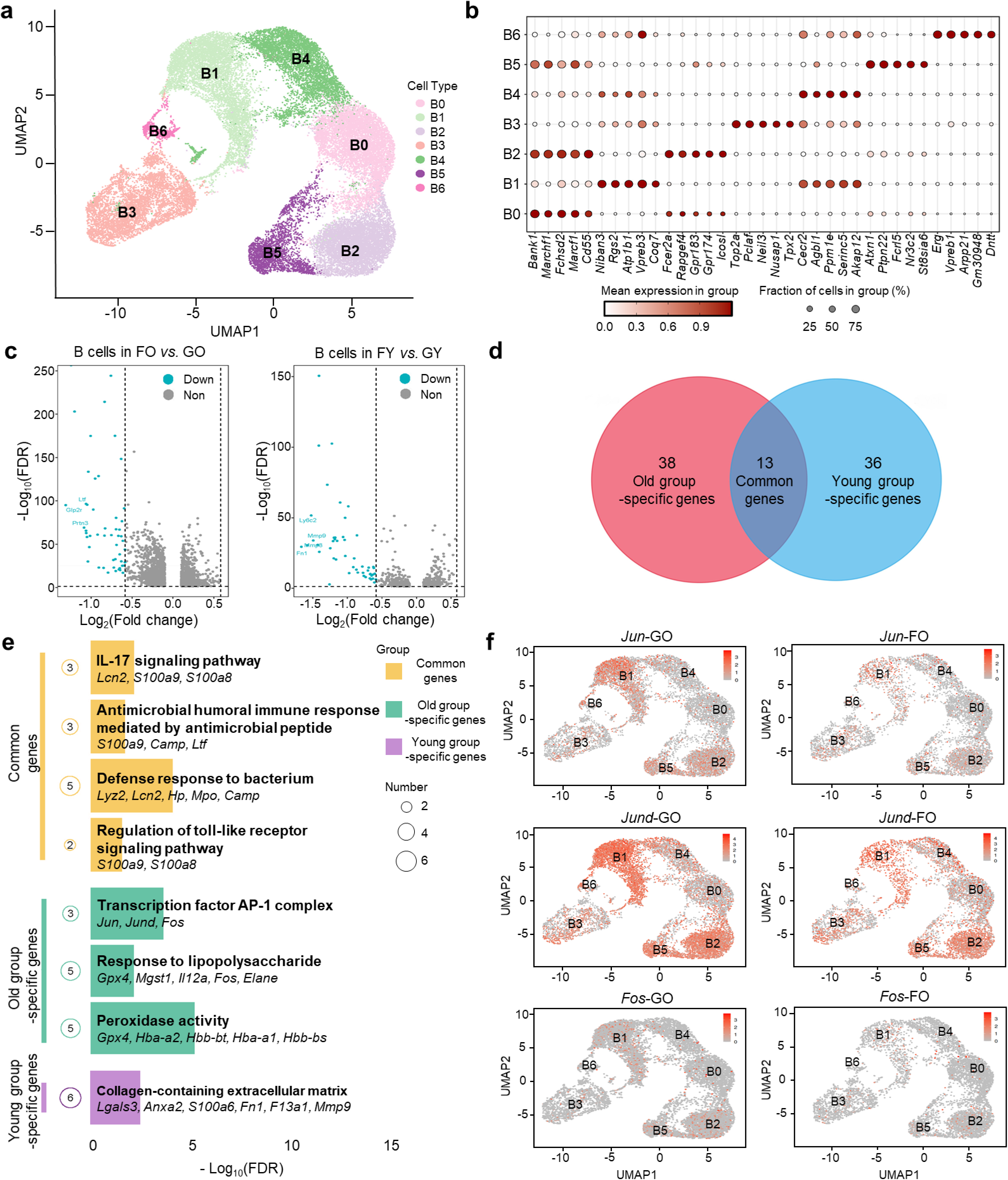
Landscape and functional features of B cells in bone marrow samples. **(a)** UMAP plot showing 34,761 B cells. **(b)** Dot plot showing the expression of canonical B cell markers used to identify B cell subtypes. **(c)** Volcano plot showing DEGs between FO and GO samples in the old group (left panel), and between FY and GY samples in the young group (right panel). Red dots represent upregulated genes, and blue dots represent downregulated genes (adjusted *p* value < 0.05 and |log_2_(fold change)| > 0.585). **(d)** Venn diagram showing overlapping downregulated DEGs in B cells from young and old groups (adjusted *p* value < 0.05, |log_2_(fold change)| > 0.585). **(e)** Functional enrichment analysis of DEGs in the young and old B cells. **(f)** UMAP plots showing the expression of Jun, Jund and Fos in aged B cells between GO and FO groups.

Spaceflight had little effect on the proportion of B cell subclusters in both young and old mice, except a decrease of cluster B1 proportion in old mice after spaceflight (Supplementary Fig 3a). However, spaceflight had a significant impact on the gene expression of B cells in both the young and old mice. We compared gene expression of FO versus GO, and FY versus GY group in B cells, and found a list of 0 upregulated and 51 downregulated genes in old mice, and 0 upregulated and 49 downregulated genes in young mice (Fig. 5c, d, adjusted *p*-value < 0.05, |log_2_(fold change)| > 0.585). Gene enrichment analysis of the 13 common downregulated genes (Fig. 5d) revealed significant enrichment in the IL-17 signaling pathway (mmu04657), antimicrobial humoral immune response mediated by antimicrobial peptides (mmu04657), defense response to bacteria (GO:0042742), and regulation of toll-like receptor signaling pathway (GO:0034121). Key genes within this common gene signature include *S100a8*, *S100a9*, *Lcn2*, *Camp*, *Ltf*, *Lyz2*, *Hp*, and *Mpo* (Fig. 5e, Supplementary Table 9). Among these, proteins encoded by *S100a8* and *S100a9* are Ca^2+^-binding proteins belonging to the S100 family, and play a crucial role in resisting microbial infection and maintaining immune homeostasis. They interact with cell surface receptors to initiate inflammatory signal transduction, induce cytokine expression and participate in the inflammatory response and immune regulation^43^. UMAP plots validated the downregulation of *S100a8* and *S100a9* in FO versus GO, and in FY versus GY group (Supplementary Fig. 3b, 3c), implying a potential impairment B cells’ capacity to respond to microbial threats, possibly contributing to increased susceptibility to infection during and after spaceflight.

Genes uniquely downregulated in old mice were enriched in pathways related to transcription factor AP-1 complex (GO:0035976), response to lipopolysaccharide (GO:0032496) and peroxidase activity (GO:0004601). The AP-1 family of transcription factors regulates a wide range of cellular processes, including cell proliferation, death, survival and differentiation^44^. In immune cells, AP-1 is activated in response to inflammatory cytokines and stress^45^. UMAP plots validates the downregulation of AP-1 transcription factors (*Jun*, *Jund*, and *Fos*), with the most pronounced changes observed in subclusters B1 and B2 (Fig. 5f), implying that B cells in old mice may have a diminished ability to initiate appropriate transcriptional programs in response to stress, potentially contributing to immune dysfunction during spaceflight.

To further understand the dynamic transcriptional changes in B cells, we performed pseudotime developmental trajectory analysis on B cell subpopulations. B3 subpopulation was defined as the starting point for trajectory analysis due to the highest potency score (Supplementary Fig. 3d). The pseudotime developmental trajectories showed only minor difference in B cell development between the spaceflight and ground control in both young and old mice (Supplementary Fig. 3e), suggesting that spaceflight has little effect on B cell development.

### Cell-cell communications

To evaluate intercellular interactions between different types of cells in spaceflight and ground control groups, we constructed a cell-cell communication network using CellChat^46,47^, based on known ligand-receptor pairs and their co-factors. We analyzed cell-cell communication characteristics separately for young and old mice, including interaction numbers, interaction strengths, and information flow.

In old mice, differential cell-cell communication analysis by comparing FO with GO group demonstrated that erythroid cells in the FO group, as information receivers, exhibited increased communication with monocytes, neutrophils, and NK-T cells (Fig. 6a, Supplementary Fig. 4a). Analysis of signaling pathways based on relative and absolute information flow revealed that pathways such as ANNEXIN, GALECTIN, VISFATIN, GRN, COMPLEMENT, and HGF were increased in the FO group, whereas the CXCL signaling pathway was decreased (Fig. 6b).

**Fig. 6.**
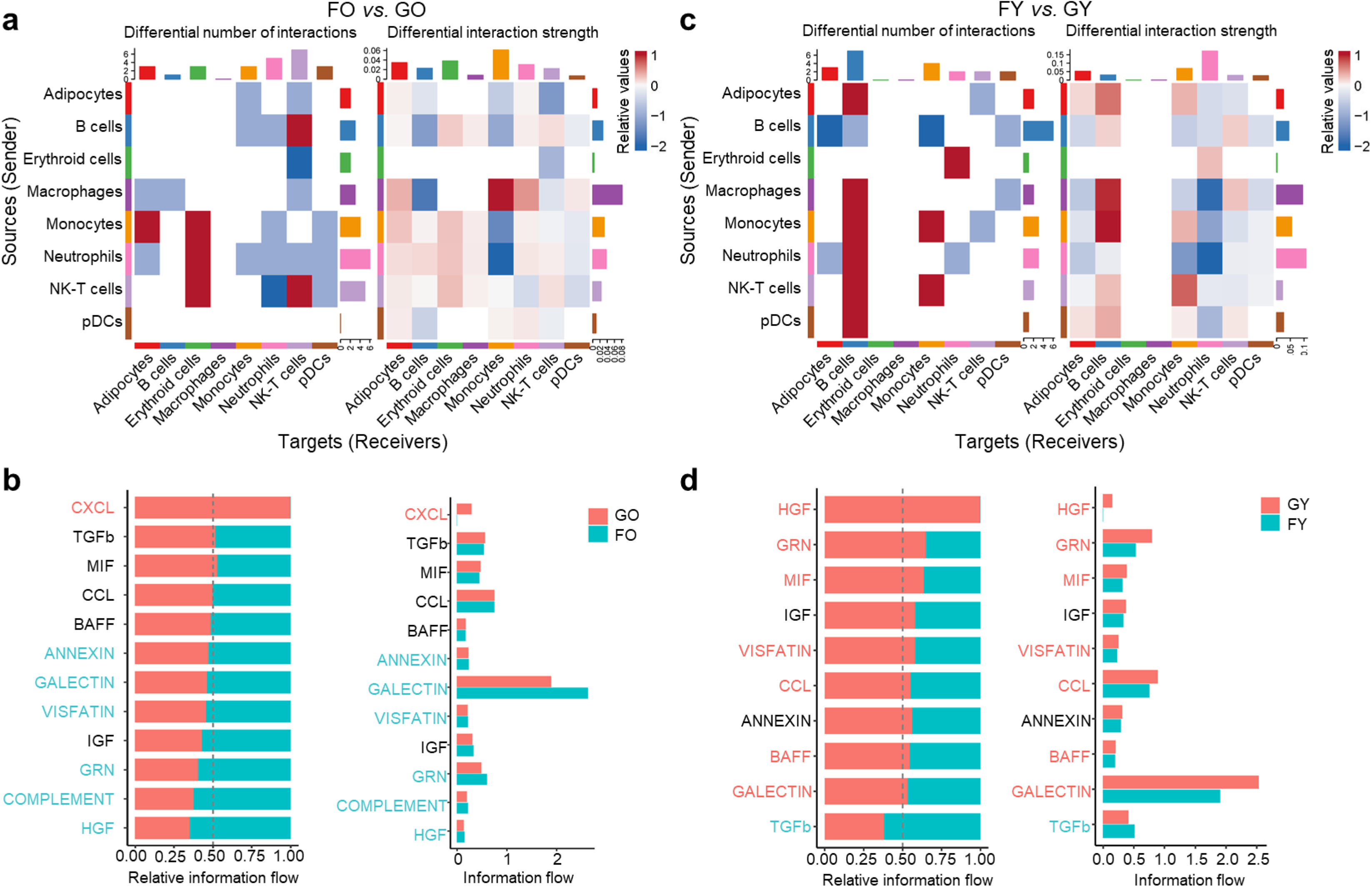
Cell-cell communication analysis across all cell types. **(a)** Heatmaps showing differential numbers (left panel) and strength (right panel) of inferred ligand-receptor interactions in the old group (FO versus GO). **(b)** Total information flow and ranking of key signaling pathways in the old group (FO versus GO). **(c)** Heatmaps showing differential numbers (left panel) and strength (right panel) of inferred ligand-receptor interactions in the young group (FY versus GY). **(d)** Total information flow and ranking of key signaling pathways in the young group (FY versus GY).

In young mice, B cells in the FY group, as information receivers, exhibited increased communication frequency and strength with multiple cell types, including adipocytes, macrophages, monocytes, NK-T cells and pDCs. However, as information senders, B cells showed decreased communication with adipocytes, monocytes and pDCs (Fig. 6c, Supplementary Fig. 4b). Analysis of signaling pathways based on relative and absolute information flow revealed that TGF-β signaling was enhanced in the FY group, whereas pathways such as HGF, GRN, MIF, VISFATIN, CCL, BAFF, and GALECTIN were significantly suppressed (Fig. 6d). Taken together, these data demonstrate that spaceflight distinctly remodels intercellular communication networks in old and young mice.

## Discussion

Spaceflight presents a unique combination of microgravity and cosmic radiation that accelerates many aging-like physiological changes, which raise concerns for astronaut health and immune resilience during prolonged missions. The bone marrow is emerging as a central nexus of spaceflight-induced aging effects, given its essential role in hematopoiesis and immune cell development^48^, and recent studies have demonstrated profound hematological stress during and after space mission. However, most space biology studies have focused on healthy young adult subjects, and it is poorly understood whether older organisms experience more severe or persistent bone marrow dysfunction after spaceflight. In addition, most prior work has largely relied on bulk measurements or ground-based analogs, which cannot fully represent the cellular heterogeneity and mechanism of bone marrow adaptation in response to spaceflight. In the present study, we utilized scRNA-seq to dissect the age-dependent effects of spaceflight on the bone marrow hematopoietic and immune system, providing a high-resolution bone marrow snapshot of the cellular landscape after a 32-day spaceflight mission followed a 24-day recovery period. Our findings reveal that spaceflight induces a state of accelerated aging in the hematopoietic system, with older mice exhibiting a profound and persistent dysregulation even after recovery. This work extends prior ground-based analogs and provide a critical high-resolution and in-vivo evidence from actual spaceflight, uncovering specific cellular vulnerabilities and molecular pathways that are exacerbated by age.

A central finding of our study is the acceleration of aging signatures within the erythroid lineage of old flight mice. While space anemia is a well-documented consequence of space travel^49^, our data provides novel mechanistic insights into the reason why recovery from space anemia is blunted in old mice. We have identified disruptions characterized by impaired erythrocyte development, inhibited mitophagy, and increased oxidative stress in the bone marrow of old mice. The suppression of mitophagy is especially significant, as this process is essential for clearing mitochondria during terminal erythroid differentiation and is a classic hallmark of cellular aging^50^. Inhibition of mitophagy suggests that aged erythroid progenitors fail to mature properly after spaceflight, leading to an accumulation of dysfunctional, ROS-producing mitochondria, which further fuels oxidative stress and cellular damage. This vicious cycle of erythroid dysfunction provides a mechanistic basis for diminished erythroid recovery in old mice after spaceflight. In contrast, erythroid cells in young mice do not show these severe changes and are more resilient, consistent with a more effective compensatory response in the young bone marrow.

Our analysis has also uncovered a persistent dysregulation of B cell populations at a single-cell resolution from actual spaceflight samples. Previous hindlimb unloading studies demonstrated a reduction in early B cell populations, and our data have confirmed a significant post-flight decrease in the overall B cell proportion, especially in older animals. At the molecular level, young and old flight groups exhibit a common downregulation of genes involved in antimicrobial humoral response, such as *S100a8* and *S100a9*, suggesting a general and lasting impairment in the innate immune capacity of B cells following spaceflight. More strikingly, we identified a unique downregulation of the AP-1 transcription factor complex (*Jun*, *Jund*, *Fos*) after spaceflight, specifically in the B cells in old mice. The suppression of AP-1, which is a critical regulator of cellular responses to stress, inflammation, and differentiation, in aged B cells, impairs the transcriptional programs for recovery, making old mice more susceptible to post-flight immune challenges.

Beyond individual lineages, spaceflight remodeled the intercellular communication network within the bone marrow, with age again as a determinant of severity. Using cell-cell communication analysis, we have found that old and young mice respond very differently in their intercellular signaling landscapes. Notably, the CXCL signaling pathway is almost completely lost in old mice after spaceflight. The CXCL axis is fundamental for the proper trafficking, retention, and function of hematopoietic stem and progenitor cells^51^, as well as for coordinating immune cell responses^52^. The disappearance of CXCL signals represents a failure in cellular crosstalk that could have cascading effects. The loss of CXCL-mediated interactions in old mice likely contributes to the inefficient reconstitution of blood cells and may partly explain the excess bone marrow fat and poor hematopoiesis. While the communication network in young mice was also altered, the changes were more of a functional shift, with TGF-β signaling becoming more prominent, again highlighting the resilience of the younger hematopoietic system.

Consistent with these signaling disturbances, we have observed changes in the bone marrow microenvironment composition that further indicate an impaired regenerative niche in old mice. One noteworthy observation is an increase in bone marrow adipocyte proportion in old mice after spaceflight compared to ground control. Bone marrow adipose tissue is known to expand with natural aging and in states of hematopoietic stress or inactivity. Our observation suggests that older bone marrow is less efficient to mobilize energy reserves for tissue regeneration. This may result from the disruption of CXCL signaling network, which tilts the balance toward adipogenesis at the expense of hematopoiesis. In essence, the aged marrow niche after spaceflight appears skewed toward a more adipose, less hematopoietically active state, which is a hallmark of marrow aging.

While our study provides unprecedented insight into cellular and molecular effects of spaceflight on bone marrow, several limitations should be acknowledged when interpreting the results. First, our analysis is based on a single recovery time point (24 days after return), which captures a snapshot of recovery but not the full temporal dynamics. A longitudinal study with multiple time points would be valuable to distinguish transient changes from permanent maladaptations and to determine how long the observed dysregulations persist. Second, as an in vivo mouse study of spaceflight, it is difficult to isolate which effects are caused by microgravity, radiation, or other space-related factors, which are intertwined during actual mission. Third, our data is collected from female mice using bone marrow from long bones (femur and humerus) rather than vertebrate. Sex and bone site differences are known to influence hematopoietic outcomes on Earth, and they could modify responses to spaceflight as well. Finally, our conclusions are based on transcriptomics and require validation with proteomic and functional assays to confirm changes at the protein and cellular activity levels. Despite these caveats, this work provides a robust single-cell atlas of the post-spaceflight bone marrow, laying the groundwork for future targeted investigations.

In conclusion, our scRNA-seq analysis provides high resolution insight into the age-dependent response of the bone marrow to spaceflight, confirming that space travel acts as a potent accelerator of hematopoietic aging. The persistent impairment of erythroid maturation, the specific failure of stress-response pathways like AP-1 in aged B cells, and the collapse of the vital CXCL communication network collectively paint a picture of decreased capacity of recovery in old mice. These findings not only deepen our understanding of spaceflight physiology but also highlight potential age-specific biomarkers and therapeutic targets, such as antioxidants to combat oxidative stress or interventions to restore CXCL signaling, to ensure the health and safety of astronauts on future long-duration missions.

## Methods

### Data acquisition

The single-cell datasets were obtained from the NASA Genelab: OSD402 (DOI: 10.26030/r127-mc55) and OSD403 (DOI:10.26030/kkf4-p733). In total, our study encompassed 148,571 cells derived from 32 samples for discovering analysis.

### Single cell RNA-seq data processing

Downstream analyses were conducted in R (version 4.4.3), primarily utilizing the Seurat^53^ R package (versions 5.0.0) along with custom scripts. The raw gene expression matrices were imported and converted into Seurat objects for subsequent analysis. To ensure high data quality, we excluded cells with low RNA content or high mitochondrial gene expression. Specifically, cells were filtered out if they had fewer than 500 RNA counts, fewer than 200 detected features, or if mitochondrial RNA content exceeded 15%. Potential doublets were identified and removed using the DoubletFinder^54^ R package (version 2.0.6) with parameter set as nExp = 7.5% for each sample.

The top 2,000 most variable genes were identified using Seurat’s “FindVariableFeatures” function. Principal Component Analysis (PCA) was subsequently performed based on these genes. To correct for batch effects across datasets, the Harmony algorithm was applied prior to clustering. Cell clustering was performed using Seurat’s “FindClusters” function. Multiple resolutions (ranging from 0.1 to 1) were tested, and the most appropriate resolution was selected based on clustering performance and biological interpretability. Then cell clusters were annotated based on the expression of known marker genes or by referencing established cell type signatures from the literature.

### Functional enrichment analysis

Differentially expressed genes (DEGs) for each cluster were identified using the “FindAllMarkers” function in the Seurat package with default parameters. Genes with an adjusted p-value < 0.05 were considered significant. A log_2_(fold change) > 0.585 was defined as upregulated, whereas a log_2_(fold change) < −0.585 indicated downregulation. We used R package clusterProfiler (version 4.14.6) to determine the enrichment of Gene Ontology (GO) and Kyoto Encyclopedia of Genes and Genomes (KEGG) pathways.

### Trajectory analysis

The CytoTRACE^55^ algorithm, an unsupervised computational framework for predicting cellular differentiation status from scRNA-seq data, was employed to assess stemness levels. CytoTRACE calculates a score for each individual cell based on the gene expression matrix, where a higher score (closer to 1) indicates greater stemness (less differentiated) and a lower score (closer to 0) indicates lower stemness (more differentiated). In this study, CytoTRACE2 R package was used to assess the differentiation status of malignant cells, B cells and erythroid cells. Visualization of differentiation trends was performed using the CytoTRACE package’s “plotCytoGenes” and “plotCytoTRACE” functions.

To reconstruct developmental trajectories, we performed pseudotime analysis using the Monocle3^56^ R package (version 1.3.7). Seurat objects were split into flight and ground control groups for separate trajectory inference. The workflow included data preprocessing, dimensionality reduction (UMAP), cell clustering by “cluster_cells” function, learning a trajectory graph by “learn_graph” function, and ordering the cells by “order_cells” function. By calculating “pseudotime,” cells were ordered along a path representing dynamic processes like differentiation.

### AUCell scoring

We used the single-cell regulatory network inference, clustering (SCENIC)^57^ R package (version 1.3.1) method, and its “AUCell” function to score the activity of 11 aging-related gene sets in each individual cell. For each cell, AUCell ranks all genes by their expression level, and calculates Area Under the Curve (AUC) to determine whether genes from a given set are enriched at the top of this ranked list.

### Cell-cell communication

We used the CellChat ^46,47^ R package (version 2.2.0) to quantitatively infer and analyze intercellular communication networks from the scRNA-seq. This workflow involves a curated database of ligand-receptor interactions to calculate communication probability between cell types based on gene expression, accounting for complex multi-subunit cofactors. The output is a communication network that can be visualized and analyzed to identify key signaling pathways, central cell types, and to compare communication patterns across different biological conditions, providing deep insights into cellular crosstalk.

### Statistics Analysis

All statistical analyses were performed using R (version 4.4.3). For group comparisons, categorical variables were analyzed with the χ² test or Fisher’s exact test, while continuous variables were assessed using the Wilcoxon rank-sum test or Kruskal-Wallis test. Pearson correlation tests were used to evaluate relationships between variables.

## Data availability

Source data for this study are publicly available in the GeneLab data repository (genelab.nasa.gov) under the Accession codes OSD-402, OSD-403. All relevant data are available from the authors.

## Code availability

All programs, packages, and functions used in this manuscript are freely available and are referred to in the “Methods” section.

## Acknowledgements

This work was supported by National Natural Science Foundation of China (Overseas Excellent Young Scientist Fund Program to H.L.; 82473126 to H.L.), Natural Science Foundation of Shandong Province (ZR2021YQ50 to H.L.; ZR2022QC212 to G.J.), Cutting Edge Development Fund of Advanced Medical Research Institute at Shandong University (GYY2023QY01 to H.L.), and Instrument Development Project of Shandong University (zy20250302 to H.L.). Haiquan Lu is a Taishan Scholar Young Talent Professor of Shandong Province and Distinguished Young Professor at Shandong University.

## Supplementary figure legends

**Fig. S1.**
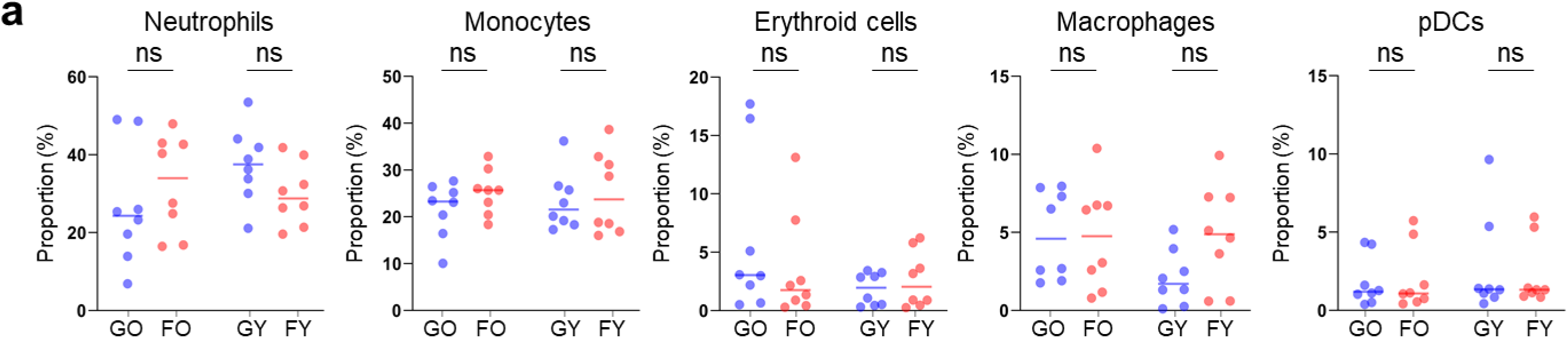
Proportional changes in major immune cell types. **(a)** Dot plots showing changes in proportion of neutrophils, monocytes, erythroid cells, macrophages and pDCs between GO and FO groups, GY and FY groups (mean ± SEM; n = 8). ns, not significant.

**Fig. S2.**
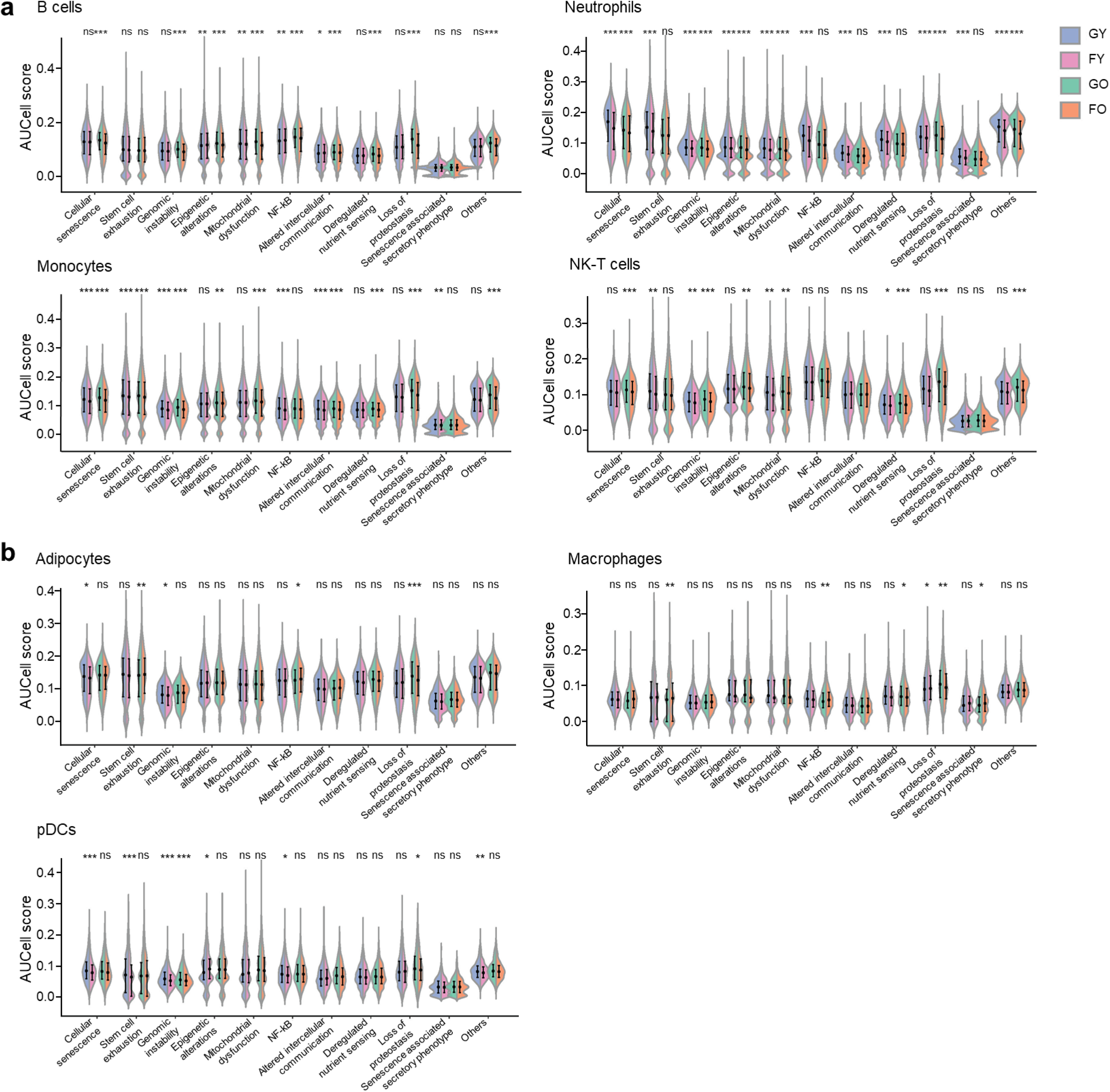
AUCell scores of aging signatures across cell types. **(a)** AUCell scores of 11 aging-related gene sets in B cells, neutrophils, monocytes and NK-T cells between GO and FO groups, and between GY and FY groups. **(b)** AUCell scores of 11 aging-related gene sets in adipocytes, macrophages and pDCs between GO and FO groups, and between GY and FY groups. \**p* < 0.05, \*\**p* < 0.01, \*\*\**p* < 0.001; ns, not significant.

**Fig. S3.**
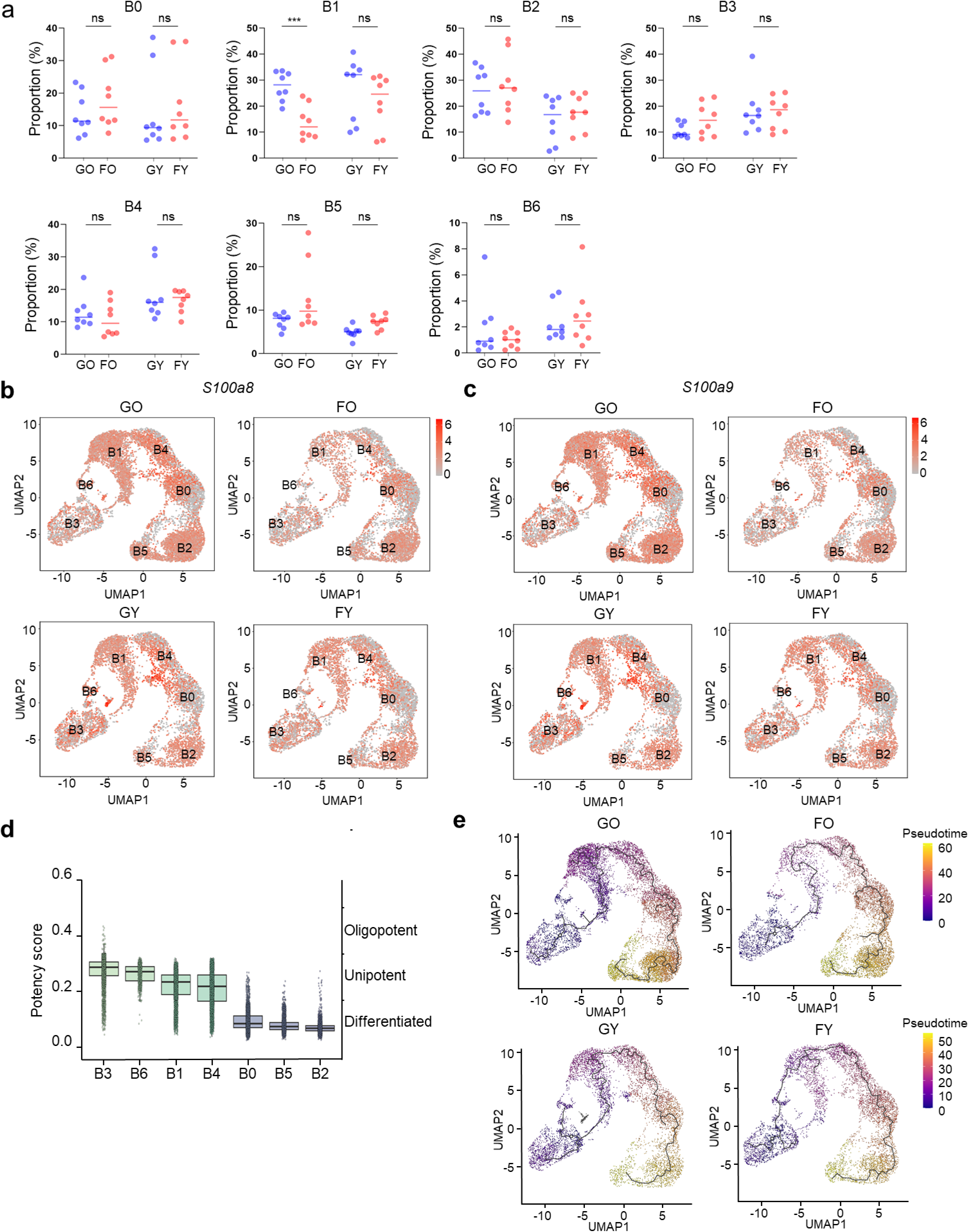
B cell subpopulation dynamics and differentiation. **(a)** Dot plots showing changes in the proportion of B0-B6 subclusters between GO and FO groups, GY and FY groups (mean ± SEM; n = 8). \*\*\**p* < 0.001; ns, not significant. **(b, c)** UMAP plots showing expression of *S100a8* and *S100a9* in B cells in GO, FO, GY and FY groups. **(d)** CytoTRACE analysis showing differentiation potential of B cell subtypes. **(e)** Trajectory analysis of erythroid cells in old (upper panel) and young (lower panel) groups under ground control and spaceflight conditions.

**Fig. S4.**
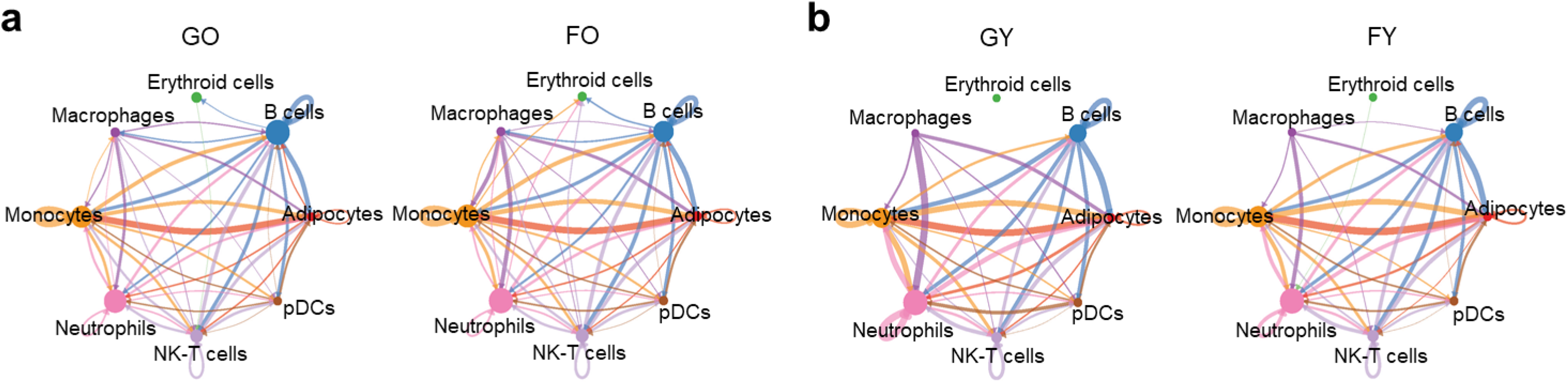
Inferred cell-cell interactions. (a, **b)** Circular plots showing the number of interactions among cell types. Line thickness represents the number of ligand-receptor interactions, and loops represent autocrine signaling.

